# Developing ERAF-AI: An Early-Stage Biotechnology Research Assessment Framework Optimized For Artificial Intelligence Integration

**DOI:** 10.1101/2025.01.08.631843

**Authors:** David Falvo, Lukas Weidener, Martin Karlsson

**Author notes:** https://www.molecule.xyz (David Falvo). https://www.bio.xyz (Lukas Weidener). https://www.coordination.network (Martin Karlsson).

## Abstract

Today, most research evaluation frameworks are designed to assess mature projects with well-defined data and clearly articulated outcomes. Yet, few, if any, are equipped to evaluate the promise of early-stage biotechnology research, which is inherently characterized by limited evidence, high uncertainty, and evolving objectives. These early-stage projects require nuanced assessments that can adapt to incomplete information, project maturity, and shifting research questions. Furthermore, these challenges are compounded by the difficulty of systematically scaling evaluations with the increasing volume of research projects. As a step toward addressing this gap, we introduce the biotechnology-oriented Early-Stage Research Assessment Framework for Artificial Intelligence (ERAF-AI), a systematic approach to evaluate research at Technology Readiness Levels (TRLs) 1 to 3 – research maturity levels where ideas are more conceptual and only preliminary evidence exists to indicate potential viability. By leveraging AI-driven methodologies and platforms such as the Coordination.Network, ERAF-AI ensures transparent, scalable, and context-sensitive evaluations that integrate research maturity classification, adaptive scoring, and strategic decision-making. Importantly, ERAF-AI aligns criteria with the unique demands of early-stage research, guiding evaluation through the 4P framework (Promote, Pause, Pivot, Perish) to inform next steps. As an initial demonstration of its potential, we apply ERAF-AI to a high-impact early-stage project, providing actionable insights and measurable improvement over conventional practices. Although ERAF-AI shows significant promise in improving the prioritization of early-stage research, further refinement, and validation across a wider range of disciplines and datasets is required to refine its scalability and adaptability. Overall, we expect this framework to serve as a valuable tool for empowering researchers to make informed decisions and to prioritize high-potential initiatives in the face of uncertainty and limited data.

## 1 Introduction

Early-stage research lays the foundation for transformative scientific and technological advancements; however, its evaluation poses significant challenges. Projects within Technology Readiness Levels (TRLs) 1 (Concept Alone) to TRL 3 (Early-Stage Evidence for Practical Use) are often conceptual or speculative and are marked by high uncertainty, limited data, and evolving hypotheses. These characteristics make early-stage research critical for innovation but are also inherently difficult to evaluate and prioritize. Traditional evaluation frameworks are not well equipped to address the unique demands of these early-stage projects, often focusing on later stages of development, where substantial experimental evidence is available. This disconnect contributes to the risk of overlooking high-potential projects, deprioritizing early-stage research, or misallocating resources. Eroom’s Law further underscores the need for improved evaluation strategies. In contrast to Moore’s Law, which predicts technological growth, Eroom’s Law highlights the declining productivity of biopharmaceutical research and development despite technological advancements (1). Addressing this stagnation requires funding high-risk, high-reward, and early-stage research. However, such efforts are hindered by inefficiencies in traditional evaluation processes, which are often time- and resource-intensive and susceptible to subjective bias.

Existing frameworks, such as the European Institute of Innovation and Technology (EIT) Pure Biotech Milestones Framework, provide structured milestone-driven guidelines for biotechnology projects (2). While these frameworks are valuable for later-stage research, they fall short of addressing the speculative and foundational nature of early-stage projects at TRLs 1–3, which often lack the extensive experimental data required to meet predefined milestones. This gap highlights the need for a framework explicitly tailored to early-stage research, incorporating flexibility to adapt to incomplete data and evolving objectives, and scalability to address the increasing volume of projects.

Recent advancements in artificial intelligence (AI) have provided a promising opportunity to transform the evaluation of early-stage research. Since its conceptual origins in the mid-20th century, AI has evolved into a powerful tool capable of solving complex problems in diverse fields (3). Applications such as PaperQA2, a frontier language model optimized for factuality, illustrate AI’s ability to enhance research workflows by performing nuanced literature search tasks, including information retrieval, summarization, and contradiction detection (4). By leveraging AI, evaluations can become more scalable, objective, and consistent, thereby addressing many of the limitations of traditional methods. However, for AI-driven evaluations to be effective, frameworks must be designed to be compatible with AI methodologies. This requires structured, machine-readable criteria, transparent decision-making processes, and the ability to integrate ethical considerations such as explainability and bias mitigation (5). Frameworks that meet these criteria can unlock the full potential of AI, enabling faster, more accurate evaluations of early-stage research while fostering transparency and reproducibility.

This manuscript introduces the *Early-Stage Research Assessment Framework for Artificial Intelligence* (ERAF-AI), specifically designed to address these challenges. ERAF-AI offers a structured, scalable, and AI-compatible approach for systematically evaluating projects at TRLs 1–3. By combining qualitative and quantitative metrics with transparent scoring and decision-making processes, ERAF-AI seeks to bridge the gap between traditional evaluation methods and the capabilities of modern AI, paving the way for more effective prioritization and funding of early-stage innovations.

### 1.1 Problem Statement

Despite the importance of early-stage research, there is a lack of evaluation frameworks specifically tailored to projects at TRLs 1–3. Existing frameworks, although effective for later-stage development, do not provide the structured, detailed criteria needed to assess the foundational and speculative nature of early-stage research. In addition, most frameworks are not designed to integrate seamlessly with AI, limiting opportunities to leverage its scalability, objectivity, and reproducibility. Addressing these gaps is critical for enabling consistent, precise, and efficient evaluation processes that support robust decision making in early-stage research.

## 2 Objectives

This article aims to develop ERAF-AI (Early-stage Research Assessment Framework for Artificial Intelligence), a framework specifically designed to evaluate early-stage research projects at Technology Readiness Levels (TRLs) 1–3. The development of ERAF-AI focuses on creating a structured methodology that can be applied systematically and consistently, enabling seamless integration with AI to enhance scalability, objectivity, and reproducibility in evaluations.

Framework Development Objectives:

- *Classification*: Establish a robust process to categorize projects into TRLs 1–3, ensuring that evaluation criteria are tailored to the specific maturity and challenges of each stage.
- *Evaluation*: Define detailed, weighted criteria for assessing key factors such as novelty, impact, and executability, structured to support integration with AI-driven tools for consistent and automated assessments.
- *Scoring and Decision-Making*: Develop a scoring system that consolidates evaluation criteria into actionable insights and guides decisions using the 4P’s framework - Promote, Pause, Pivot, or Perish.

By addressing the gaps in existing evaluation frameworks and prioritizing compatibility with AI methodologies, ERAF-AI seeks to create a transparent, reproducible, and scalable approach for assessing the potential of early-stage research projects, providing clarity and rigor to decision-making processes.

## 3 Methodology

ERAF-AI was developed to systematically evaluate early-stage research projects, focusing on classification, evaluation, scoring, and decision-making. The framework was designed to ensure compatibility with both human evaluators and AI systems, emphasizing structured criteria, weighted scoring, and transparent decision-making processes tailored to TRLs 1–3.

### 3.1 Initial Framework Development

ERAF-AI was developed to provide a structured approach for evaluating early-stage research projects. It combines qualitative and quantitative metrics through a scoring system designed to ensure systematic, transparent, and repeatable evaluations. Each criterion is assigned a weighted importance value reflecting its significance at a specific TRL. Projects are scored on a scale of 1 (excellent) to 5 (poor), with scores multiplied by their respective weights to ultimately produce weighted scores. Using this approach provides a comprehensive view of a project’s potential, while also prioritizing critical factors relevant to the corresponding project maturity level.

ERAF-AI employs the 1-5 scoring scale for its balance between granularity and simplicity, enabling nuanced evaluations without adding unnecessary complexity. Binary scoring systems lack sufficient differentiation, and confidence intervals or percentage-based scoring can complicate aggregation and prioritization. By aligning with established practices in both academic and industry contexts, ERAF-AI ensures usability while enabling adaptability across diverse early-stage projects.

ERAF-AI comprises three main steps:

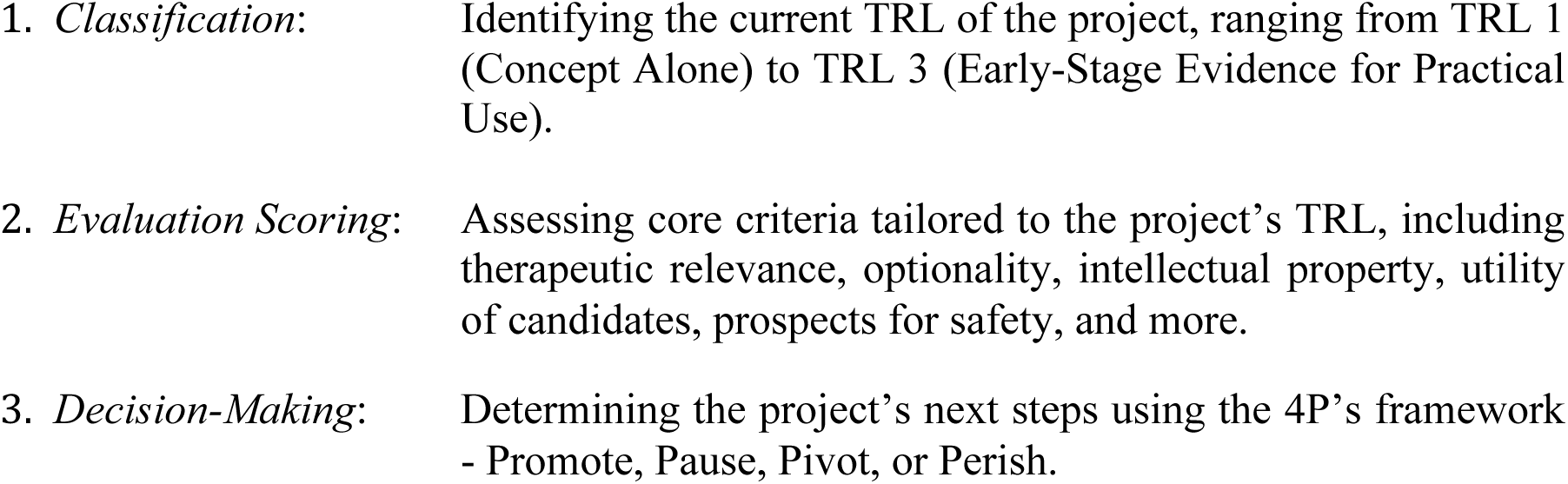

The ERAF-AI framework was grounded in a comprehensive review of existing methodologies, including established frameworks in biotechnology and pharmaceutical innovation. A database search was conducted in PubMed, Scopus, and Web of Science, using keywords such as ‘research evaluation frameworks,’ ‘biotechnology assessment,’ and ‘TRL-based evaluation.’ The search focused on publications from the last 15 years to ensure relevance to current practices and advancements. Records were screened for their applicability to TRLs 1-3, with an emphasis on frameworks addressing scalability, decision-making, and practical evaluation. The database search was further supplemented by an online search using the same keywords through a general search engine (Google.com) to capture relevant unpublished or non-commercially published documents and additional resources. Moreover, insights and resources were obtained through consultations with experts in the biotechnology field, leveraging professional networks to identify key frameworks and methodologies that may not be readily accessible in the indexed databases. The ERAF-AI framework represents a significant development in evaluating early-stage biotechnology research ventures, particularly when compared with established valuation methods. Traditional approaches, such as the Risk-Adjusted Net Present Value method (rNPV), the Discounted Cash Flow Method (DCF), Comparable Company Analysis (CCA), and the Venture Capital (VC) method rely heavily on financial metrics, market-based comparables, or speculative projections, making them ill-suited for pre-revenue, early-stage biotech startups. While most effective for later-stage ventures, these methods struggle to capture the complexities and uncertainties inherent in TRLs 1–3, where robust financial data are absent and progress hinges on scientific and technological milestones. When compared to qualitative methods, such as the Berkus Method and the Scorecard Valuation Method, ERAF-AI offers superior granularity and adaptability to the biotech sector. The Berkus Method, with its simplified monetary assignment to broad categories, lacks the depth needed to evaluate nuanced biotech projects tied to evidence-based milestones and intellectual property rather than more generic startup metrics (e.g., prototype completion). The Scorecard Valuation Method is more nuanced, utilizing weighted criteria such as team strength and market opportunity, but it remains static and subjective, heavily dependent on local market comparables, and often ill-defined reasoning. Additionally, the EIT Pure Biotech Milestones Framework, which is robust in defining biotechnological milestones, lacks a structured scoring system or weighting mechanism, which makes systematic cross-project comparisons difficult. Another identified and relevant framework, Jack Scannell’s Drug Project Guidance, includes broad and overlapping categories, but does not offer explicit methods for decision-making or assessing scalability (6).

ERAF-AI distinguishes itself through its AI-driven design, which ensures consistency and traceable reasoning in evaluations, critical improvements over the subjectivity of traditional frameworks. By prioritizing stage-appropriate scientific, technological, and operational metrics, ERAF-AI offers tailored evaluations of biotech, focusing on factors such as therapeutic relevance, scalability, and regulatory readiness. The dynamic, weighted scoring system adapts to evolving priorities and uncertainties, ensuring forward-looking, justifiable assessments. Additionally, ERAF-AI incorporates iterative re-evaluation, enabling continuous refinement as projects progress to maturity. This structured emphasis on transparency, scalability, and adaptability allows the ERAF-AI to bridge the gap between theoretical evaluation models and practical applications. With the integration of AI, not only is there enhanced decision-making through the 4P framework (Promote, Pause, Pivot, Perish), but there is also an opportunity to address the biotechnology industry’s pressing need for actionable, data-driven, and context-sensitive assessment frameworks, offering a new benchmark in early-stage research evaluation.

The development process also involved iterative refinement through collaboration with two additional experts specializing in biotechnology and project evaluation. These experts provided critical feedback on the framework’s structure, evaluation criteria, and scoring methodology, ensuring alignment with both academic rigor and industry best practices. Weighted importance values were carefully calibrated during this process, informed by insights from the systematic review and expert feedback. These values reflect the evolving priorities of projects across TRLs 1–3, and ensure that ERAF-AI provides a fair and adaptable assessment framework for early-stage innovations.

### 3.2 Artificial Intelligence Integration

ERAF-AI was intentionally designed to integrate seamlessly with generative AI models, such as ChatGPT, to enhance the evaluation process of early-stage research projects. This integration enables automated, consistent, and scalable assessments while preserving transparency and usability for human evaluators. The development of ERAF-AI was guided by the necessity to create a framework that AI systems can interpret, apply, and refine autonomously or in collaboration with human oversight. In keeping with this objective, future iterations can incorporate language agents to engage with the human-in-the-loop cycles of conjecture and criticism. By analyzing extant scientific literature to highlight areas of contention, contextualizing hypotheses within the broader research landscape, and offering hypothesis-specific inferences, agents can offer personalized evaluations for human researchers to help spur more fruitful investigations at scale.

To ensure AI usability while retaining the framework’s rigor and adaptability, the following principles were embedded into ERAF-AI’s design:

- *Machine-Readable Evaluation Framework*: The scoring system, classification criteria, and decision-making steps were structured in formats that can be directly processed by AI systems. This includes defined workflows and logical pathways that allow generative AI to consistently replicate human decision-making (7).
- *Standardized and Modular Criteria*: Evaluation criteria for each TRL were standardized and modularized, enabling AI systems to apply TRL-specific criteria dynamically. This modularity allows the framework to evolve as AI capabilities improve without disrupting its foundational structure.
- *AI-Driven Scoring Logic*: The scoring system was designed to enable generative AI models to calculate weighted scores based on input data. Each scoring criterion includes structured prompts that guide AI tools in analyzing inputs, evaluating evidence, and assigning scores objectively (8).
- *Traceable Decision Pathways*: The 4P’s framework (Promote, Pause, Pivot, Perish) was adapted to include clear, traceable logic pathways. This enables AI systems to provide transparent justifications for their recommendations, ensuring accountability and interpretability (9).

ERAF-AI leverages the unique strengths of generative AI by embedding the following functionalities:

- *Prompt Optimization for TRL Classification and Evaluation*: AI systems are guided to classify projects by asking predefined questions aligned with TRL criteria. These prompts were crafted to ensure AI models interpret project data in line with the objectives of the ERAF-AI (10).
- *Automated Decision-Support Tools*: Generative AI systems are used to automate parts of the scoring and decision-making process, particularly for data-rich evaluations such as intellectual property analysis or safety assessments.
- *Natural Language Processing (NLP) for Insights*: AI models analyze unstructured data, such as project descriptions or scientific literature, to extract insights relevant to ERAF-AI’s core evaluation criteria.

To ensure functionality and adherence to best practices for AI compatibility, the initial framework was reviewed by an expert specializing in AI applications. This review focused on validating the framework’s machine-readability, modularity, and traceability, ensuring that it aligns with industry standards for AI usability (11, 12). The expert also provided feedback on integrating ethical considerations, such as transparency, bias mitigation, and explainability, into the design.

### 3.3 The Coordination.Network: AI-Augmented Research

The Coordination.Network is a decentralized, open-source platform that accelerates scientific discovery by coordinating researchers, domain experts, and funding bodies through AI-enhanced collaboration. Complementing AI-driven evaluation systems such as ERAF-AI, it promotes scalability, transparency, and accountability by enabling participants to develop automated workflows or “skills.” These skills encapsulate expert methodologies for tasks, such as proposal evaluation, literature synthesis, and experimental design, ensuring thorough oversight while scaling knowledge across distributed teams. By embedding expertise into AI-compatible processes, the platform streamlines the transition from theoretical research workflows to practical applications. Its progressively decentralized infrastructure and blockchain-based smart contracts maintain transparent governance, secure data sharing, and trustless collaboration. A local-first approach grants researchers full control over self-hosted instances, whereas a centrally hosted version supports early adopters. As the network expands, independently hosted instances are integrated into a unified ecosystem guided by AI-assisted consensus building and dynamic analysis of project metrics and contributor feedback to inform decisions on research evaluation, funding allocation, and collaborative governance.

Structured workflows guide AI systems and collaborators through predefined methodologies to ensure consistency, transparency, and rigorous oversight. Iterative feedback loops incorporate skill versioning with ongoing community review, maintaining accuracy, and fostering the continual improvement of AI-generated insights. Currently, verification depends on trusted contributors, but future enhancements will leverage advanced AI models to automate and refine this process. Open APIs facilitate seamless integration with external computational tools, databases, and machine-learning pipelines, supporting data exchange and hypothesis generation. Researchers can customize and combine skills for specific needs, enabling near-real-time responsiveness in dynamic environments. A token-based incentive system, alongside a reputation framework, rewards high-quality research contributions and rigorous peer review, whereas human oversight preserves adaptability and accountability in governance and decision-making. Positioned at the frontier of AI-augmented research, the Coordination.Network is evolving in step with advancements in decentralized computing and human-machine collaboration. Autonomous agents are poised to propose and validate hypotheses by synthesizing data from diverse repositories, thereby increasing the efficiency of experimental methods while preserving human oversight. The open-source foundation supports prospective integration with emerging human-AI interfaces, potentially including brain-computer technology. Over time, AI will move beyond analytic support to autonomously generate and adapt research skills with human experts ensuring the alignment of strategic objectives. By combining AI-based evaluation (ERAF-AI) and AI-enhanced coordination (Coordination.Network), this ecosystem provides a robust infrastructure for research execution, from hypothesis generation to funding decisions, while upholding transparency, scalability, and academic rigor.

### 3.4 Applying ERAF-AI

To preliminarily validate ERAF-AI’s functionality and compatibility with AI systems, the Coordination.Network was chosen as the testing platform. The Coordination.Network was selected for its transparency in decision-making, ensuring clear rationale and traceability for every output generated, as well as its support for interchangeable large language models (LLMs), such as OpenAI’s ChatGPT. This interoperability allows for robust experimentation with different generative AI systems while maintaining consistent evaluation standards.

For this validation, publicly available research information was sourced using the NIH RePORTER database.

The following selection criteria were applied to identify relevant projects:

- *Fiscal Year*: 2020–2024
- *NIH Spending Category*: Biotechnology
- *Activity Codes*: R01, R03, R21, SBIR/STTR
- *Award Type*: New

These criteria yielded 16 total results. From this list, the project with the highest total cost was selected for evaluation. This approach ensured that the chosen project was highly funded, reflecting a significant investment in early-stage research and aligning with ERAF-AI’s focus on impactful assessments. The full project data can be found in the Appendix (A).

## 4 Results

The ERAF-AI framework was developed to address the challenges of evaluating early-stage research by providing a structured, transparent, and actionable methodology. The framework consists of four main components: Classification, Evaluation, Scoring, and Decision-Making. Each component builds upon the previous one, creating a comprehensive approach that aligns project assessments with their stage of development and readiness for advancement.

### 4.1 Classification

The first step of the ERAF-AI framework ensures that projects are assessed based on their maturity and progress within the TRL system. Accurate classification is crucial for tailoring evaluation criteria to the specific needs and challenges of each developmental stage. Projects are classified into one of the following levels:

**TRL 1: Concept (Theoretical Stage):**

At this stage, research focuses purely on conceptual or theoretical work, with no experimental evidence or practical applications.

***Key Criteria for TRL 1:***

**1. Evidence of Research Activities:**

- No experimental research or testing conducted.
- Documentation includes theoretical models, literature reviews, or white papers without empirical data.
**2. Nature of Work:**

- The project is entirely focused on theoretical or conceptual ideas.
- No proof-of-concept experiments are present.

**3. Scientific Hypothesis:**

- A biologically plausible hypothesis may exist but lacks any experimental validation.
- Hypothesis strength is evaluated based on alignment with existing scientific knowledge.

***Examples of TRL 1:***

**•** A proposed novel drug target based on theoretical insights.
**•** Computational simulations predicting likely efficacy of a molecule, with no experimental confirmation.

**TRL 2: Speculative Research Conducted (Early Feasibility Stage):**

At TRL 2, initial feasibility studies or speculative research are performed to test the theory. Experiments may be limited in scope, speculative, or inconclusive.

***Key Criteria for TRL 2:***

1. Evidence of Experimental Research:

- Early feasibility or speculative research (e.g., initial in vitro experiments, computational simulations with early validation) has been conducted.
- Experimental design may lack robustness but provides preliminary insights.
2. Support for Hypothesis:

- Early-stage experiments support the hypothesis but are not definitive.
- Results may indicate potential mechanisms of action or initial feasibility.
3. Emerging Evidence:

- Emerging experimental evidence validating the concept or mechanism of action is present, even if incomplete or preliminary.

***Examples of TRL 2:***

- Initial in vitro studies showing that a compound binds to a target of interest.
- Preliminary screening assays identifying potential relationships between target proteins and an *in vitro* phenotype, with no follow-up validation studies conducted.

***TRL 3: Early-Stage Evidence for Practical Use (Foundational Research)***:

At this stage, there should be foundational research supporting its practical use, with experiments confirming early feasibility in relevant preclinical disease models (e.g., rodents, patient-derived organoids).

***Key Criteria for TRL 3:***

1. **Experimental Evidence in Models:**

- Experiments conducted in relevant in vivo or in vitro models (e.g., animal disease models, organoids) confirm early feasibility.
- Evidence must demonstrate potential applicability to real-world use cases.
2. **Efficacy Demonstration:**

- Early-stage efficacy is shown in preclinical models (e.g., reduction in tumor size in a rodent model).
- Results support the hypothesis with measurable outcomes.
3. **Preliminary Safety and Scalability Considerations:**

- Indications of safety (e.g., absence of major toxicities in preclinical tests).
- Early consideration of development factors like Good Manufacturing Practice (GMP) or Chemistry, Manufacturing, and Controls (CMC).

***Examples of TRL 3:***

- Preclinical efficacy data from in vivo studies showing disease amelioration.
- Preliminary safety testing showing low toxicity in preclinical models.

This classification system is essential for ensuring fairness and relevance in project evaluations. By aligning evaluation priorities with the project’s maturity, ERAF-AI allows for consistent, stage-appropriate assessments that guide early-stage research toward meaningful milestones. ERAF-AI’s classification system was designed to leverage AI for enhanced accuracy and scalability. Machine-readable formats were applied to TRL definitions and criteria, enabling AI models to automate classification by interpreting input data through standardized prompts. Additionally, AI tools such as generative models can dynamically assess project descriptions, assigning TRL classifications based on key project features and identified gaps. By integrating AI into this stage, ERAF-AI ensures that classification is consistent, objective, and repeatable, reducing variability and bias inherent in manual assessments. AI’s ability to continuously refine classification logic through iterative feedback also enhances the system’s adaptability over time.

### 4.2 Evaluation Scoring

Building on the classification step, which identifies the appropriate TRL for each project, the evaluation process focuses on assessing projects based on their specific TRL. By tailoring evaluation criteria to the challenges and opportunities unique to each TRL, ERAF-AI ensures that projects are assessed fairly and effectively. This stage represents the core of the framework, providing actionable insights aligned with the project’s developmental maturity. The following sections detail the evaluation approach for each TRL.

#### 4.2.1 ERAF-AI Technology Readiness Level 1

ERAF-AI TRL 1 evaluates research at a theoretical level, where concepts and mechanisms are proposed but not yet experimentally validated. At this stage, the bar for success focuses on the novelty, theoretical soundness, and potential impact of the idea. Since no experiments have been conducted, the evaluation emphasizes theoretical grounding and innovation, with a lower stringency for feasibility and data requirements. However, projects must demonstrate strong potential through scientifically plausible mechanisms, biological relevance, and a solid intellectual property position.

**Evaluation Objectives for TRL 1**

1. Therapeutic Relevance of Mechanism of Action (Weighted Importance: 60%)

- *Rationale*: The proposed mechanism of action should be scientifically plausible and biologically relevant, even if it remains hypothetical. This ensures that the concept aligns with a clear therapeutic need and provides a strong foundation for further exploration.
- *Questions to Consider*:

- Is the proposed mechanism scientifically plausible and logically consistent?
- Does the concept target a biologically relevant disease mechanism or unmet need?
- Does the concept address a significant unmet medical need in a sizable patient population?
- *Scoring Guidelines (1 to 5, lower is better)*:

- 1-2: The mechanism is strongly biologically relevant and logically consistent.
- 3: The mechanism has some biological plausibility but presents notable uncertainties.
- 4-5: The mechanism is speculative or lacks coherence and logical consistency.
2. Therapeutic Optionality (Weighted Importance: 15%)

- *Rationale*: The mechanism should demonstrate flexibility by potentially addressing multiple therapeutic areas or indications. This optionality increases the concept’s robustness and broadens its potential applications.
- *Questions to Consider*:

- Could this mechanism potentially address multiple therapeutic areas or indications?
- Are there potential commercial applications for the mechanism across multiple indications or markets?
- *Scoring Guidelines (1 to 5, lower is better)*:

- 1-2: The mechanism has strong potential for multiple applications.
- 3: The mechanism shows limited but feasible flexibility for additional uses.
- 4-5: The mechanism has narrow applicability or questionable potential for expansion.
3. Intellectual Property (Weighted Importance: 25%)

- *Rationale*: At this stage, novelty is crucial. The concept must demonstrate clear potential for patentability while minimizing risks of prior art or competition. Strong intellectual property ensures that the concept has a defensible position as it progresses.
- *Questions to Consider*:

- Does the concept represent a novel approach that can be patented?
- Are there significant risks of prior art or existing competitive solutions?
- Does the concept demonstrate a non-obvious inventive step, or would a person with ordinary skill in the field readily arrive at the same solution through routine methods or straightforward experimentation?
4. *Scoring Guidelines (1 to 5, lower is better)*:

- 1-2: The concept has clear potential for strong IP protection with minimal risks of prior art.
- 3: The concept is novel but presents some potential risks related to prior art or competition.
- 4-5: The concept lacks novelty or has significant risks of prior art, making IP protection weak.

ERAF-AI TRL 1 differentiates itself from the EIT Pure Biotech Milestones Framework and Jack Scannell’s Drug Project Guidance by introducing specific weighted criteria tailored to theoretical concepts (2, 6). While the EIT framework focuses on milestone-driven progress, such as identifying unmet needs and conducting initial patent assessments, ERAF-AI quantitatively evaluates novelty, biological relevance, and IP strength. The explicit weighting ensures that these foundational aspects are prioritized, addressing a gap in EIT’s milestone-driven approach.

**Table 1:** TRL 1 - Concept (Theoretical Stage)

| Criterion | Weighted Importance (%) | Score (1-5) | Weighted Score |
| --- | --- | --- | --- |
| Therapeutic Relevance of Mechanism | 60% |  |  |
| Therapeutic Optionality | 15% |  |  |
| Intellectual Property | 25% |  |  |

Unlike Scannell’s guidance, which employs broad and static categories, ERAF-AI incorporates dynamic, iterative evaluations based on emerging data. This adaptability allows for reassessment as new information becomes available, ensuring that theoretical concepts are evaluated comprehensively and fairly. ERAF-AI’s tailored approach makes it particularly suited for early-stage projects requiring rigorous evaluation of theoretical potential.

AI tools are used to enhance the consistency and scalability of TRL 1 evaluations. Generative AI models can automate scoring by interpreting project descriptions through predefined prompts aligned with TRL 1 criteria. This ensures objective and reproducible evaluations, particularly for novelty and intellectual property assessments. Additionally, AI models dynamically adapt to evolving project data, allowing iterative refinement of evaluations based on new insights. These AI-driven enhancements ensure that ERAF-AI integrates cutting-edge automation without compromising its methodological rigor.

#### 4.2.2 ERAF-AI Technology Readiness Level 2

ERAF-AI TRL 2 evaluates research at an early feasibility stage, where speculative research has been conducted to validate initial hypotheses and explore the practical application of proposed mechanisms. At this stage, the balance between theoretical soundness and experimental evidence becomes more critical. Projects are expected to provide early-stage experimental support for their hypotheses, even if the data remains speculative. The goal is to assess whether early feasibility efforts validate the proposed mechanism while identifying potential risks or areas for improvement.

**Evaluation Objectives for TRL 2**

1. Therapeutic Relevance of Mechanism of Action (Weighted Importance: 45%)
2. *Rationale*: Early experimental results should begin to validate the theoretical hypothesis established in TRL These results must demonstrate biological relevance and reinforce the feasibility of the proposed target.
3. *Questions to Consider*:

**–** Do the early experimental results support the original hypothesis?

**–** Does the data reinforce the biological relevance of the proposed target?

- *Scoring Guidelines (1 to 5, lower is better)*:

**–** 1-2: Experimental results strongly support the hypothesis with high biological relevance.

**–** 3: Limited experimental evidence with some gaps in biological relevance.

**–** 4-5: Experimental results are insufficient or conflicting, undermining confidence in the hypothesis.

1. Therapeutic Optionality (Weighted Importance: 5%)
2. *Rationale*: Early findings may reveal new applications or pathways for the proposed mechanism, increasing the concept’s flexibility and potential impact.
3. *Questions to Consider*:

**–** Are there emerging findings suggesting alternative therapeutic applications?

**–** Can this mechanism be applied to other indications or pathways?

- *Scoring Guidelines (1 to 5, lower is better)*:

**–** 1-2: Strong indications of broader applications based on early findings.

**–** 3: Limited but viable potential for additional applications.

**–** 4-5: No clear evidence of therapeutic optionality.

1. Intellectual Property (Weighted Importance: 15%)
2. *Rationale*: Early feasibility results should strengthen the project’s novelty and support potential patent filings. IP strategy at this stage may involve formalizing patents or mitigating prior art risks.
3. *Questions to Consider*:

**–** Can early feasibility results strengthen IP filings?

**–** Is there a clear pathway to formalize patent applications?

- *Scoring Guidelines (1 to 5, lower is better)*:

**–** 1-2: Early data strongly supports patentability and novel approaches.

**–** 3: Some uncertainties about IP protection or competition.

**–** 4-5: Significant challenges to securing strong IP, such as prior art risks.

1. Utility of Candidates (Weighted Importance: 20%)
2. *Rationale*: At TRL 2, potential therapeutic candidates should begin to emerge. These candidates, while still speculative, must demonstrate preliminary viability in achieving therapeutic objectives.
3. *Questions to Consider*:

**–** Are potential therapeutic candidates emerging from the early research?

**–** How viable are these candidates in achieving the desired therapeutic effects

**–** Do the emerging drug candidates have a competitive advantage over existing or emerging therapies?

- *Scoring Guidelines (1 to 5, lower is better)*:

**–** 1-2: Promising candidates identified with strong rationale for their selection.
**–** 3: Limited evidence supporting candidate viability.
**–** 4-5: High uncertainty with no clear candidates identified.
- Prospects for Safety (Weighted Importance: 15%)

- *Rationale*: Early indications of safety or toxicity risks should emerge at this stage, even if data is speculative. Identifying potential safety concerns early reduces downstream risks.
- *Questions to Consider*:

**–** Are there early signs of potential safety concerns, such as toxicity or off-target effects?

**–** Do the emerging safety and toxicity profiles compare favorably to those of existing treatments on the market?

- *Scoring Guidelines (1 to 5, lower is better)*:

**–** 1-2: No significant safety concerns identified in early research.

**–** 3: Some manageable safety concerns.

**–** 4-5: Early data shows significant safety risks that could hinder progress.

ERAF-AI TRL 2 differentiates itself from the EIT Pure Biotech Milestones Framework and Jack Scannell’s Drug Project Guidance by introducing weighted assessments tailored to early feasibility (2, 6). While the EIT framework focuses on exploratory milestones like defining biotechnological platforms, ERAF-AI explicitly evaluates Utility of Candidates (20%) and Prospects for Safety (15%), ensuring that emerging therapeutic candidates and early safety signals are systematically addressed.

In contrast to Scannell’s broader and overlapping categories, such as “safety prospects” and “clinical feasibility,” ERAF-AI offers granular, weighted criteria that emphasize feasibility and risk management at the early stages. By focusing on specific indicators such as candidate viability and safety risks, ERAF-AI provides a structured and actionable framework for addressing critical uncertainties during the transition from conceptual validation to experimental feasibility.

**Table 2:** TRL 2 - Speculative Research Conducted (Early Feasibility Stage)

| Criterion | Weighted Importance (%) | Score (1-5) | Weighted Score |
| --- | --- | --- | --- |
| Therapeutic Relevance of Mechanism | 45% |  |  |
| Therapeutic Optionality | 5% |  |  |
| Intellectual Property | 15% |  |  |
| Utility of Candidates | 20% |  |  |
| Prospects for Safety | 15% |  |  |

To enhance consistency and scalability, AI tools were integrated into TRL 2 evaluations. Generative AI models automate scoring for criteria like Utility of Candidates and Prospects for Safety, using structured prompts and predefined workflows to interpret experimental data. AI can dynamically adapt to new findings, allowing iterative refinement of assessments while maintaining traceability and transparency. By leveraging these capabilities, ERAFAI ensures robust and reproducible evaluations that guide early-stage projects effectively toward TRL 3.

#### 4.2.3 ERAF-AI Technology Readiness Level 3

Building on the foundation established in the classification step, the evaluation process for ERAF-AI TRL 3 focuses on advanced feasibility, where experimental evidence substantiates the practical application of proposed mechanisms. At this stage, foundational research must provide strong evidence of efficacy, safety, and scalability, marking a critical transition from speculative feasibility to preclinical readiness. With increased expectations for validation, projects face higher scrutiny, and gaps in feasibility or experimental data weigh more heavily on evaluations.

ERAF-AI TRL 3 builds upon the objectives assessed in TRL 1 and TRL 2 (therapeutic relevance, therapeutic optionality, intellectual property, utility of candidates, and safety) while introducing four additional focus areas critical for preclinical readiness: prospects for GMP/CMC for IND filing, prospects for clinical development, commercial potential, and organization and team fit.

**Evaluation Objectives for TRL 3**

1. Therapeutic Relevance of Mechanism of Action (Weighted Importance: 20%)

- *Rationale*: Research at this stage must provide clear experimental evidence that the proposed mechanism has strong therapeutic relevance.
- *Questions to Consider*:

**–** Does the research demonstrate that the mechanism can be exploited therapeutically?
**–** Are the experimental results consistent with achieving therapeutic relevance?
- Scoring Guidelines (1 to 5, lower is better):

**–** 1–2: Strong experimental validation supporting therapeutic relevance.
**–** 3: Moderate evidence with areas needing further confirmation.
**–** 4–5: Insufficient or conflicting evidence undermining therapeutic relevance.
2. Therapeutic Optionality (Weighted Importance: 5%)

- *Rationale*: Clear indications should emerge at this stage for expanding the mechanism into additional therapeutic areas, enhancing its flexibility and impact.
- *Questions to Consider*:

**–** Are there clear indications for expanding into other disease areas?
**–** Could the mechanism be adapted to additional therapeutic pathways?
- Scoring Guidelines (1 to 5, lower is better):

**–** 1–2: Strong evidence of broader applicability.
**–** 3: Limited but feasible potential for expansion.
**–** 4–5: Narrow applicability with little or no therapeutic optionality.
3. Intellectual Property (Weighted Importance: 15%)

- *Rationale*: Foundational data should confirm the concept’s novelty and feasibility for securing robust IP protection.
- *Questions to Consider*:

**–** Does the data reinforce the novelty of the concept?
**–** Is there a robust strategy in place for IP filings or licensing opportunities?
- Scoring Guidelines (1 to 5, lower is better):

**–** 1–2: Strong IP position supported by foundational data.
**–** 3: Some risks to IP strength or novelty but manageable.
**–** 4–5: Significant barriers to obtaining or defending IP.
4. Utility of Candidates (Weighted Importance: 20%)

- *Rationale*: Candidates at this stage should demonstrate early efficacy in disease models, providing confidence in their potential for further development.
- *Questions to Consider*:

**–** Do the candidates show clear evidence of efficacy in preclinical disease models?
**–** Are the candidates viable for achieving the desired therapeutic effects?
- Scoring Guidelines (1 to 5, lower is better):

**–** 1–2: Strong evidence of efficacy in preclinical models.
**–** 3: Partial efficacy but requires further validation.
**–** 4–5: Lack of efficacy or unreliable preclinical results.
5. Prospects for Safety (Weighted Importance: 15%)

- *Rationale*: Safety profiles should emerge from early in vivo studies, addressing toxicity risks and supporting further development.
- *Questions to Consider*:

**–** Are there early indications of safety and low toxicity from preclinical studies?
**–** Are risks manageable, or do they present significant challenges?
- Scoring Guidelines (1 to 5, lower is better):

**–** 1–2: No significant safety concerns identified.
**–** 3: Moderate risks but manageable with mitigation.
**–** 4–5: Significant safety concerns or toxicity risks.
6. Prospects for GMP/CMC for IND Filing (Weighted Importance: 5%)

- *Rationale*: Manufacturing and scalability readiness should be assessed to ensure candidates can be produced under GMP conditions for IND filing.
- *Questions to Consider*:

**–** Is there a clear pathway for manufacturing and scaling the candidates?
**–** Are there significant barriers to GMP or CMC compliance?
- Scoring Guidelines (1 to 5, lower is better):

**–** 1–2: Strong prospects for GMP readiness and scalability.
**–** 3: Manageable challenges to scaling.
**–** 4–5: Significant barriers to GMP readiness.
7. Prospects for Clinical Development (Weighted Importance: 10%)

- *Rationale*: This step evaluates the feasibility of transitioning to clinical trials, including regulatory, logistical, and resource requirements.
- *Questions to Consider*:

**–** Is there a clear pathway for transitioning to clinical trials?
**–** Are there significant regulatory or logistical hurdles?
- Scoring Guidelines (1 to 5, lower is better):

**–** 1–2: Clear and feasible pathway for clinical development.
**–** 3: Moderate challenges but solvable.
**–** 4–5: Major roadblocks to clinical trials.
8. Commercial Potential (Weighted Importance: 5%)

- *Rationale*: Projects at this stage should demonstrate market demand and competitive potential to support development.
- *Questions to Consider*:

**–** Are there clear commercial applications?
**–** What is the estimated market size and potential revenue for the therapeutic candidate?
**–** What is the competitive landscape and potential market share?
**–** What are the potential barriers to entry for new competitors?
- Scoring Guidelines (1 to 5, lower is better):

**–** 1–2: High commercial viability with strong market potential.
**–** 3: Moderate commercial potential with some challenges.
**–** 4–5: Limited commercial feasibility or intense competition.
9. Organization and Team Fit (Weighted Importance: 5%)

- Rationale: The organization and team must demonstrate the capability to execute the next stages of development, including clinical and commercial phases.
- Questions to Consider:

**–** Does the team have the expertise and resources to advance the project?
**–** Are there gaps in expertise or resources that could hinder progress?
- Scoring Guidelines (1 to 5, lower is better):

**–** 1–2: Strong team and resources to execute plans effectively.
**–** 3: Team has gaps but can address them.
**–** 4–5: Team lacks key expertise or resources for advancement.

ERAF-AI TRL 3 distinguishes itself from the EIT Pure Biotech Milestones Framework and Jack Scannell’s Drug Project Guidance by addressing key gaps in preclinical readiness, operational planning, and decision-making through its granular and weighted criteria.

The EIT framework primarily emphasizes milestone-driven progress, such as initiating CMC development and defining target product profiles (TPPs). However, it lacks a structured, weighted approach for systematically evaluating preclinical feasibility, GMP readiness, and scalability. For example, while EIT highlights milestones like “initial CMC development,” ERAF-AI TRL 3 assigns specific weights (5%) to Prospects for GMP/CMC for IND Filing, ensuring that manufacturability is explicitly assessed as a critical standalone criterion. Furthermore, EIT’s broad focus on product profiles is refined in ERAF-AI by distinct evaluations of Commercial Potential (5%) and Utility of Candidates (20%), offering a more precise and actionable assessment of market readiness and therapeutic viability. These weighted criteria in ERAF-AI provide a structured framework for decision-making, enabling systematic project comparisons and informed prioritization.

**Table 3:** TRL 3 - Early-Stage Evidence for Practical Use (Foundational Research)

| Criterion | Weighted Importance (%) | Score (1-5) | Weighted Score |
| --- | --- | --- | --- |
| Therapeutic Relevance of Mechanism | 20% |  |  |
| Therapeutic Optionality | 5% |  |  |
| Intellectual Property | 15% |  |  |
| Utility of Candidates | 20% |  |  |
| Prospects for Safety | 15% |  |  |
| Prospects for GMP/CMC for IND Filing | 5% |  |  |
| Prospects for Clinical Development | 10% |  |  |
| Commercial Potential | 5% |  |  |
| Organization and Team Fit | 5% |  |  |

In contrast, Jack Scannell’s Drug Project Guidance focuses on broader categories such as clinical feasibility, safety prospects, and therapeutic relevance but lacks detailed subcategories tailored to preclinical readiness (6). For instance, while Scannell includes general safety assessments, ERAF-AI assigns a detailed weight (15%) to Prospects for Safety, explicitly addressing early toxicity risks and safety signals emerging from preclinical models. This ensures a systematic approach to safety evaluation, which Scannell’s framework does not provide at the same level of granularity.

Additionally, Scannell’s reliance on subjective confidence scoring introduces variability and reduces transparency in evaluations. ERAF-AI counters this limitation by providing explicit scoring guidelines for all criteria, such as Prospects for Clinical Development (10%) and Organization and Team Fit (5%), ensuring that project feasibility and team capabilities are systematically and transparently evaluated. Moreover, Scannell’s framework omits GMP readiness as a specific evaluation category, whereas ERAF-AI explicitly incorporates it, addressing critical considerations for manufacturability and scale-up potential early in the evaluation process.

ERAF-AI’s unique contribution lies in its ability to integrate preclinical readiness, regulatory considerations, and strategic alignment into a single, cohesive framework. By embedding operational and strategic factors such as GMP readiness, commercial potential, and team capacity into weighted evaluations, ERAF-AI offers actionable insights that are often absent or underdeveloped in EIT and Scannell’s frameworks. Unlike the static, milestone-driven approach of EIT or the broader categorizations in Scannell, ERAF-AI employs specific weighted criteria tailored to the needs of TRL 3, ensuring adaptability and rigor. The framework dynamically adjusts its focus to include priorities like clinical development feasibility (10%) and commercial viability (5%), providing a comprehensive evaluation of a project’s readiness to advance to clinical stages.

The scoring process in the ERAF-AI framework synthesizes the evaluation results into an overall success score that reflects the novelty, impact, and executability of a project. This step enables the effective ranking of projects within the same TRL, ensuring that decision-making prioritizes the most promising candidates while maintaining transparency and consistency across evaluations. The scoring approach is designed to adapt to the evolving priorities at each TRL, emphasizing novelty and impact at earlier stages and placing greater weight on executability at more advanced stages. The primary goal of the scoring process is to derive a comprehensive success score for each project, reflecting its potential to advance through development.

The scoring system integrates structured prompts and machine-readable formats, ensuring compatibility with AI systems for automated calculations. This enhances scalability and objectivity, enabling consistent application of scoring logic across diverse projects. ERAF-AI’s scoring approach ensures that projects are systematically evaluated, prioritizing those with the greatest potential for advancement while aligning with the framework’s emphasis on transparency and adaptability.

#### 4.2.4 TRL-Speci/ic Weighting

**TRL 1:** At the conceptual stage, novelty and impact carry more weight, as these are the primary drivers of success. Executability has lower stringency, given the early nature of the research.

**TRL 2:** At the feasibility stage, the balance shifts slightly, with executability beginning to gain importance. Utility of Candidates and safety prospects start to play a larger role in defining the project’s potential.

**TRL 3:** At the preclinical readiness stage, executability carries the greatest weight. This reflects the need for strong candidates, robust safety data, and a capable team to ensure successful advancement. Impact is now closely tied to real-world efficacy, safety, and development potential.

The final success score is calculated as the average of the three factors: Novelty, Impact, and Executability. Scores are rated on a scale from 1 to 5, with lower scores indicating greater potential for success. An overview of the final scoring is shown in Table 5.

### 4.3 Decision-Making

The final step of the ERAF-AI framework translates evaluation scores into actionable decisions, ensuring that resources are allocated effectively, and projects are advanced, refined, or discontinued based on their potential and alignment with TRL-specific goals. Decision-making integrates both human judgment and AI capabilities to reassess a project’s TRL classification as new data becomes available, applying criteria to determine the most appropriate next steps.

The primary goal of the decision-making process is to make clear, informed choices about a project’s trajectory. This includes evaluating whether the project should advance to the next TRL, require further development, pivot to a new direction, or be abandoned entirely. By leveraging AI for automated reassessment and consistency, ERAF-AI ensures scalable and objective decision-making while maintaining transparency.

#### 4.3.1 The 4P Framework

- **Promote:** Should the project advance to the next TRL based on strong alignment with the current TRL criteria and evidence of readiness?
- **Pause:** Are there gaps in data, feasibility, or evidence that necessitate further research or refinement before promotion?
- **Pivot:** Should resources be reallocated, or should the project pivot to a new direction based on identified limitations or opportunities?
- **Perish:** Should the project be discontinued due to fundamental flaws, disproven hypotheses, or a lack of feasibility?

Projects are assessed for readiness to advance to the next TRL based on their final success score, with specific thresholds applied to determine their readiness. An overview is shown in Table 6

**Table 4:** Final Scoring Range.

| Final Score Range | Interpretation |
| --- | --- |
| 1-2 | High potential for success |
| 3 | Moderate potential; requires refinement |
| 4-5 | High risk with low likelihood of success |

**Table 5:** Overview of the 4P Framework for Decision-Making.

| Decision | Criteria |
| --- | --- |
| Promote | Strong alignment with TRL stage; scores primarily in the $1.0 \leq \text{Score} < 2.0$ range |
| Pause | Moderate alignment with TRL stage; some gaps in feasibility or evidence require addressing $2.0 \leq \text{Score} < 3.0$ |
| Pivot | Misalignment with TRL stage but potential for adaptation through reallocation of resources or altered strategy |
| Perish | Fundamental misalignment or lack of feasibility; scores primarily in the $4.0 \leq \text{Score} \leq 5.0$ range |

**Table 6:** TRL Decision Thresholds.

| Score Range | Category | Readiness Level | Decision |
| --- | --- | --- | --- |
| $1.0 \leq \text{Score} < 2.0$ | Excellent Project | High Readiness | Promote to the next TRL with confidence |
| $2.0 \leq \text{Score} < 3.0$ | Promising Project | Moderate Readiness | Pause to address gaps or advance conditionally upon resolving specific challenges |
| $3.0 \leq \text{Score} < 4.0$ | Moderate Project | Low Readiness | Significant refinement or foundational work required before promotion |
| $4.0 \leq \text{Score} \leq 5.0$ | High Risk / Low Potential | Not Ready | Strongly consider project termination or substantial restructuring |

#### 4.3.2 Thresholds for TRL Promotion

**Excellent Project (1.0 ≤ Score *<* 2.0):**

These projects show high novelty, strong early-stage evidence, and excellent potential for development. They fully align with the criteria of their current TRL and are well-positioned to progress to the next stage.

**Promising Project (2.0 ≤ Score *<* 3.0):**

These projects demonstrate significant promise but require refinement or further validation to strengthen their alignment with TRL criteria. Advancement may be conditional on addressing specific gaps.

**Moderate Project (3.0 ≤ Score *<* 4.0):**

These projects face substantial risks or misalignments with the current TRL stage. Additional foundational work or significant refinement is required before they can progress.

**High Risk / Low Potential (4.0 ≤ Score ≤ 5.0):**

These projects are unlikely to succeed without major changes or breakthroughs. They are strong candidates for termination or substantial restructuring.

#### 4.3.3 Dynamic Reassessment of TRL Classification

As new data becomes available, a project’s TRL classification should be reassessed to ensure the evaluation criteria remain appropriate and reflective of its current maturity. The rationale and criteria outlined in Step 1: Classification are revisited to determine whether promotion to a higher TRL is warranted.

This iterative approach ensures flexibility and responsiveness to evolving project circumstances. AI systems embedded within ERAF-AI play a critical role in this process, automating reassessments and providing consistent, unbiased recommendations based on updated evaluation data. The ability to dynamically adjust TRL classifications enhances the framework’s adaptability and ensures that project trajectories align with both emerging data and strategic objectives. An overview of the decision-making and promotion process of the ERAF-AI is shown in Table 7.

**Table 7:** Case Study: Manually Organized Data Following AI Evaluation

| Criterion | Weighted Importance (%) | Score (1-5) | Weighted Score |
| --- | --- | --- | --- |
| Therapeutic Relevance of Mechanism | 20% | 1.5 | 0.3 |
| Therapeutic Optionality | 5% | 1.5 | 0.075 |
| Intellectual Property | 15% | 1.5 | 0.225 |
| Utility of Candidates | 20% | 1.5 | 0.30 |
| Prospects for Safety | 15% | 1.5 | 0.225 |
| Prospects for GMP/CMC for IND Filing | 5% | 3 | 0.15 |
| Prospects for Clinical Development | 10% | 2 | 0.20 |
| Commercial Potential | 5% | 1.5 | 0.075 |
| Organization and Team Fit | 5% | 1 | 0.5 |

## 5 Implementing ERAF-AI: A Case Study

To preliminarily validate ERAF-AI in a practical setting, the Coordination.Network was selected as the implementation platform, leveraging its transparency and flexibility in integrating predefined research project data. The identified project, detailed in Appendix A, provided a relevant use case for ERAF-AI’s evaluation capabilities.

### 5.1 Overview of Selected Project

- **Project Title:** Rapid response for pandemics: single cell sequencing and deep learning to predict antibody sequences against an emerging antigen.
- **Total Funding:** $1,851,627
- **Public Health Relevance Statement:** The readiness for a future viral pandemic will critically depend on being able to accurately predict, entirely using computers, neutralizing antibody structures directly from the virus’s protein structure. Vaccines and therapeutics can then be manufactured and tested with unprecedented rapidity. We propose to make this goal a reality by engaging the advantages of single-cell genomic deep sequencing and the latest advances in deep machine learning.

The selected project aligns well with ERAF-AI’s emphasis on early-stage research evaluation, focusing on novelty, impact, and executability.

For each of the four steps of the ERAF-AI framework, classification, evaluation, scoring, and decision-making, the corresponding data utilized within the Coordination.Network for AI-based implementation is detailed, along with the exact prompts employed to facilitate automated processing and decision-making.

### 5.2 Step 1: Classification

The objective of this step was to determine the appropriate Technology Readiness Level (TRL) of the selected research project (Appendix A) using the ERAF-AI framework in conjunction with the Coordination.Network as the AI tool.

#### 5.2.1 Data Input

- Complete research project information
- Classification details from the result section

#### 5.2.2 Prompt 1

Begin by identifying the current TRL of the project - ranging from Concept Alone (TRL 1) to Early-Stage Evidence for Practical Use (TRL 3).

#### 5.2.3 Response

To determine the current Technology Readiness Level (TRL) of the project titled “Rapid response for pandemics: single cell sequencing and deep learning to predict antibody sequences against an emerging antigen,” an assessment based on the stages of TRL 1 to TRL 3 is required:

- **TRL 1 (Concept Alone - Theoretical Stage):** Research is purely theoretical with no experimentation or proof of concept. This does not apply, as the project description includes experimental tasks.
- **TRL 2 (Speculative Research Conducted - Early Feasibility Stage):** Some early feasibility or speculative research is performed to validate the theory. The project involves developing deep learning architectures and utilizing them to discover associations; this signifies some movement beyond theoretical work.
- **TRL 3 (Early-Stage Evidence for Practical Use - Foundational Research):** Foundational research with experiments confirming early feasibility. The project’s use of LIBRA-seq to rapidly identify and generate candidate immunoglobulin coding sequences, along with validation via synthetic antibodies and phage-display systems, aligns with early-stage practical applications.

#### 5.2.4 Conclusion

Based on the outlined project tasks, including generating training sets, validating predictions, and applying foundational principles in biological engineering, the project likely aligns with TRL 3. It displays early-stage evidence supporting practical applications through structured research and experimentation.

### 5.3 Step 2: Evaluation Scoring

The objective of this step was to evaluate the selected research project (Appendix A) using the ERAF-AI framework for the assigned TRL (TRL 3).

#### 5.3.1 Data Input

- Complete research project information
- Evaluation details from the result section
- Prompt 1 response (TRL classification)

#### 5.3.2 Prompt 2

Apply the relevant evaluation criteria tailored to the project’s TRL. Each criterion is weighted based on its importance at that stage. Use the provided worksheet to systematically evaluate and score the project. If the project is deemed between TRLs, use the lower-level evaluation criteria.

#### 5.3.3 Response

Based on the provided project details and its current stage, the evaluation has been conducted using the TRL 3 criteria, as the project aligns with TRL 3:

- Therapeutic Relevance of Mechanism of Action

– **Weighted Importance:** 20%
**– Questions to Consider:** Does the research provide clear evidence that the mechanism can be exploited therapeutically?
**– Evaluation:** The project uses foundational research with LIBRA-seq and validation via synthetic antibodies and phage-display systems, demonstrating clear early-stage experimental usage.
**– Score:** 1-2 (Assumed high performance based on description)
- Therapeutic Optionality

– **Weighted Importance:** 5%
**– Questions to Consider:** Are there clear indications for expanding into other disease areas?
**– Evaluation:** Potential for broader applications beyond SARS-CoV-2, including non-infectious or chronic diseases.
– **Score:** 1-2
- Intellectual Property

– **Weighted Importance:** 15%
**– Questions to Consider:** Is there a robust IP strategy in place?
**– Evaluation:** The project describes a novel computational approach with unique single-cell sequencing, likely to have strong IP potential.
– **Score:** 1-2
- Utility of Candidates

– **Weighted Importance:** 20%
**– Questions to Consider:** Do the drug candidates show early efficacy in disease models?
**– Evaluation:** The predicted antibodies are validated experimentally showing practical applicability.
– **Score:** 1-2
- Prospects for Safety

– **Weighted Importance:** 15%
**– Questions to Consider:** Is there evidence for safety and low toxicity?
**– Evaluation:** With potential testing through phage-display systems safety aspects can be explored early.
– **Score:** 1-2
- Prospects for GMP / CMC for IND Filing

– **Weighted Importance:** 5%
**– Questions to Consider:** Can candidates be produced at scale under GMP conditions?
**– Evaluation:** Early-stage; production and scale-up might not be assessed yet but would be integral after proof of concept.
– **Score:** 3
- **Prospects for Clinical Development**

– **Weighted Importance:** 10%
**– Questions to Consider:** What are the next steps for clinical trials?
**– Evaluation:** The development will pave paths for trials, particularly in vaccine enhancement against viral pandemics.
– **Score:** 2
- **Commercial Potential**

– **Weighted Importance:** 5%
**– Questions to Consider:** How competitive is the market?
**– Evaluation:** The project likely shows high commercialization potential due to its innovative approach to preventing pandemics.
– **Score:** 1-2
- Organization and Team Fit

– **Weighted Importance:** 5%
**– Questions to Consider:** Does the team have the expertise and resources?
**– Evaluation:** Multi-institutional team with diverse expertise across computer science, immunology, and molecular biology; very strong fit.
**–** Score: **1**

5.3.4 **Conclusion**

The framework confirms the project is well-aligned with TRL 3, supported by early practical evidence and strong alignment for advancement into practical applications.

#### 5.3.5 Manually Organized Data Following AI Evaluation

Following the AI-based evaluation of the research project, the provided data is summarized in the table below. Each criterion was assessed according to its weighted importance and assigned a score based on the evaluation findings.

The total weighted score, derived from the aggregated values across all criteria, **is 1.60**. This result reflects a high readiness level and strong alignment with the expectations for TRL 3.

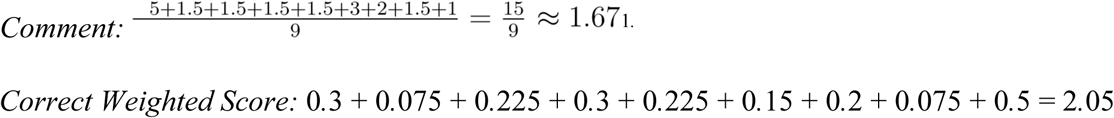

### 5.4 Step 4: Decision-Making

The objective of this step was to determine the appropriate next steps for the selected research project (Appendix A) based on its final weighted score and alignment with the decision-making criteria of the ERAF-AI framework for TRL 3.

#### 5.4.1 Data Input

- Complete research project information
- Decision-making details from the result section
- Prompt 3 response (scoring)

#### 5.4.2 Prompt 4

Decide the fate of the project (4P’s – Promote, Pause, Pivot, or Perish) based on its overall score and alignment with strategic goals.

#### 5.4.3 Response

The project’s current success score of 1.53 falls into the range for an **‘Excellent Project’** (1.0

≤ Score *<* 2.0), indicating that it shows high novelty, strong early-stage evidence, and excellent potential for development. This aligns well with the criteria for promotion to a higher Technology Readiness Level (TRL).

#### 5.4.4 Outcome

**Promote –** The project should advance to the next TRL level due to its high readiness and alignment with strategic goals. The team and the foundational framework established seem capable of supporting further advancements, and the project’s objectives align with urgent public health needs such as pandemic preparedness, leading to a strategic fit for promotion.

## 6 Discussion

The Early-Stage Research Assessment Framework for Artificial Intelligence (ERAF-AI) offers a novel solution for evaluating early-stage research projects at TRLs 1–3. By leveraging AI, ERAF-AI enhances scalability, transparency, and objectivity, addressing the limitations of traditional frameworks such as the EIT Pure Biotech Milestones Framework, which often lacks specificity and flexibility for early-stage evaluations (2). Through its modular structure, classification, evaluation, scoring, and decision-making, ERAF-AI aligns the assessment criteria with the unique challenges of early-stage research projects. The application of the framework in a case study, where the research project was classified as TRL 3, demonstrates its ability to systematically prioritize evaluation criteria and adapt to developmental maturity. The AI-classified TRL of 3 aligns with the authors’ manual evaluation using both ERAF-AI and the EIT framework. The selection of the highest-cost project for evaluation serves as an indicative factor because higher funding levels often correlate with more advanced research stages. While this does not constitute definitive proof of the project’s TRL classification, it provides a strong indication of its alignment with TRL 3, supported by foundational evidence and early feasibility. This congruence between AI-driven and manual evaluations reinforces the capacity of ERAF-AI to systematically prioritize evaluation criteria based on developmental maturity.

The ERAF-AI framework draws inspiration from traditional models, such as the EIT Pure Biotech Milestones Framework and Scannell’s Drug Project Guidance, while introducing targeted refinements to address the unique challenges of early-stage research (2, 6). Similar to the EIT framework, ERAF-AI employs a stage-gated development process using TRLs, aligning the evaluation criteria with project maturity to provide clear expectations at each stage. Moreover, ERAF-AI shares the multidimensional evaluation focus of both the EIT and Scannell frameworks by assessing the therapeutic relevance, intellectual property, safety, and commercial viability. However, ERAF-AI diverges in key areas to enhance its applicability to early-stage research projects. The TRL-based classification system provides a granular and context-specific method for mapping project maturity that is particularly relevant for academic and exploratory research. Additionally, the dynamic weighting of criteria ensures that novelty is emphasized at earlier stages, whereas feasibility and scalability gain prominence at later TRLs. The integration of scoring metrics, such as novelty, impact, and executability, further simplifies decision-making, offering actionable insights that are uniquely suited to the uncertainties of early-stage research.

The case study results underscore the importance of modular and weighted evaluation criteria, which prioritize therapeutic relevance, intellectual property, and optionality at TRL 3. Specific findings from the case study demonstrated strong alignment with TRL 3 attributes, including robust therapeutic relevance and intellectual property potential. However, challenges in areas such as GMP/CMC readiness have been identified, highlighting the capacity of ERAF-AI to identify developmental bottlenecks. These results affirm the utility of the framework in de-risking early-stage research by focusing on the critical feasibility and scalability factors. The case study also revealed domain-specific challenges, such as reliance on sufficient training datasets for AI tools, particularly in niche research areas. While the Coordination.Network’s ability to integrate interchangeable LLMs, addressed these challenges to some extent, broader access to high-quality datasets remains essential for robust implementation across disciplines.

As AI continues to redefine how scientific research is conducted and evaluated, tools such as FutureHouse’s PaperQA2 present a compelling opportunity to further enhance the capabilities of ERAF-AI (13) . PaperQA2, a state-of-the-art AI agent for high-accuracy retrieval augmented generation (RAG) from scientific documents, outperforms PhD and Postdoc-level biologists in scientific reviews, offering significant potential for addressing persistent challenges in handling large-scale literature retrieval, summarization, and contradiction detection. Thus, the integration of PaperQA2 with ERAF-AI and the Coordination.Network could enable deeper and more nuanced evaluations while streamlining the extraction of actionable insights. With the enhanced ability to process large amounts of unstructured scientific data, opportunities emerge for bridging information gaps in ways that prioritize relevant hypotheses and ultimately enable the accelerated and more systematic evaluation of research ventures without sacrificing accuracy. By incorporating tools such as PaperQA2 with ERAF-AI and the Coordination.Network, current processes can be refined in ways that lay the groundwork for a more dynamic and adaptive evaluation environment, one that evolves and scales with the growing needs of early-stage research.

The integration of AI enables scalable and objective evaluations while reducing human bias. Generative AI systems, supported by platforms such as the Coordination.Network, efficiently interpret data and apply criteria across large datasets, addressing challenges posed by the variability and uncertainty of early-stage research. For instance, the ability of AI to process unstructured data such as abstracts and protocols into structured evaluation scores highlights its potential for transformation (14). However, this reliance on AI introduces risks of over-reliance on automated outputs, particularly when the quality of the training datasets is insufficient or unrepresentative (15). While feedback loops allow for the continuous refinement of AI outputs with human expertise, this hybrid model still faces limitations in fully mitigating automation biases and ensuring interpretability across diverse applications

(16). ERAF-AI promotes transparency and accountability through machine-readable frameworks and traceable decision-making pathways. However, challenges persist, including equitable access to AI tools and addressing the biases embedded within the training data. Furthermore, maintaining meaningful human oversight is not just desirable, but essential to safeguard trust and prevent the framework from inadvertently reinforcing existing biases or errors.

Further consideration pertains to a thorough evaluation of intellectual property. Accurately determining novelty and non-obviousness requires exhaustive analysis of patent libraries, which can be as extensive as, or even larger than, scientific databases. While ERAF-AI can offer a preliminary assessment by referencing easily accessible information, the question of obviousness, whether a person with ordinary skill in the field could arrive at the same invention through straightforward experimentation, demands nuanced human judgment. In scenarios in which an existing technology is adapted for a new but closely related application, AI alone may not reliably discern this subtlety. Therefore, ERAF-AI should include explicit disclaimers clarifying that its findings do not substitute for legal counsel or a comprehensive patent search. As digital archives become more systematically curated and widely accessible, AI-driven algorithms could play a more prominent role in evaluating patentability, especially for initial screening. However, the impact of AI-driven IP assessments is likely to grow as models are **1)** continuously trained on available patent literature and **2)** taught how to discern, to some degree of probability, that a human of skill in a field would find a novel idea in that field obvious. If and when achieved, IP-literate AI models will provide new opportunities for more thorough and efficient patent searches.

An additional direction for future research is to validate ERAF-AI by using larger and more diverse datasets to determine whether AI can detect correlations or patterns that may elude human evaluators. For example, testing all 16 candidate projects in the NIH dataset and examining whether ERAF-AI’s final scores align with each project’s funding level could reveal new insights into funding decisions and highlight potential outliers, such as high-scoring yet underfunded proposals. These discrepancies might warrant re-evaluation and redirect resources to overlooked projects with high potential. Moreover, the impact of specific training data on AI-driven assessments should be examined, as domain-focused or expert-augmented data can alter how ERAF-AI ranks project novelty, feasibility, or impact.

### 6.1 Limitations

Although ERAF-AI integrates AI to enhance objectivity, scalability, and reproducibility, several limitations merit attention. First, the reliance on AI-driven methodologies introduces challenges related to data quality. Early-stage research often involves sparse or incomplete datasets, which can lead to biased or suboptimal AI output. If training datasets are unrepresentative or overly narrow, AI systems may fail to generalize effectively, potentially skewing evaluations and decision-making. The algorithmic biases inherent in training data can also propagate through AI-driven processes, inadvertently reinforcing existing inequalities or inaccuracies (17). Additionally, while feedback loops with human evaluators mitigate some of these risks, they do not entirely eliminate the possibility of over-reliance on automated outputs, particularly when evaluators lack technical expertise to critically appraise AI-generated results. Finally, as generative AI evolves, the risk of ‘black-box’ decision-making increases, and the rationale behind AI outputs becomes opaque, posing challenges to interpretability and trust (18). Addressing these limitations will require ongoing efforts to refine the training datasets, improve transparency, and ensure robust human oversight.

## 7 Conclusion

This study introduces ERAF-AI (Early-stage Research Assessment Framework for Artificial Intelligence) as a novel tool designed to address the complexities of evaluating early-stage research projects at TRLs 1–3. By leveraging AI methodologies, ERAF-AI provides a structured, scalable, and objective evaluation framework that addresses gaps in traditional evaluation systems. Its integration with the Coordination.Network highlights the potential for modularity, transparency, and adaptability in real-world applications. The case study validation demonstrates the capacity of ERAF-AI to identify key areas of strength and developmental bottlenecks in early-stage research projects, offering actionable insights for de-risking innovation pipelines. Despite these contributions, the limitations associated with the data quality, algorithmic biases, and interpretability underscore the need for further refinement and validation. Future research should focus on expanding the framework’s applicability to diverse disciplines, incorporating ethical safeguards, and iterating AI-driven methodologies to enhance reliability and trust. By addressing these areas, ERAF-AI can serve as a pivotal tool for fostering innovation in early-stage research.

## Acknowledgements

The authors express their gratitude to everyone who contributed to the creation of this research. In particular, we extend our thanks to Logan Bishop-Currey, Lutz Kummer, James Brodie, Sean Brennan, and Tyler Quigley for their valuable feedback, suggestions, and reviews.

## Appendix A Research project Information

Link: https://reporter.nih.gov/search/8MA1eUw8vEaefDvWi-lzyA/project-details/10274223

**Title:** Rapid response for pandemics: single cell sequencing and deep learning to predict antibody sequences against an emerging antigen

`o` **Project Number:** 1R01AI169543-01
`o` **Former Number:** 1R01OD031531-01
`o` **Contact PI/Project Leader:** RAY, ANIMESH
`o` **Awardee Organization:** KECK GRADUATE INST OF APPLIED LIFE SCIS
`o` **Fiscal year:** 2021
`o` **Total funding:** $1,851,627

**Abstract:** One of the “holy grails” in immunology is to be able to directly predict tight-binding variable chain antibody sequences in silico against foreign or non-self ‘antigenic’ proteins. Immunoglobulin chain rearrangement can potentially encode approximately 1016 different variants of antibody heavy and light chain sequences. However, only a small fraction of the sequence space is generally accessed for evolving antibodies against foreign proteins. The computational challenge is to go from a model of the structure of an antigen to predicting a set of antibody chain sequences that can bind tightly to the antigen. If solved, it might be possible to move in less than 24 hours from the first cryo-electron-microscopic structure of a novel viral protein to advance a set of potent antibody-like molecular candidates for testing. Towards solving this problem, this project aims to develop a deep learning architecture that will take as input thermodynamic, quantum mechanical (density functional), and local structure-based network topographical features of the antigens and their cognate antibodies, and will output their respective binding affinity constants. We will design a generative adversarial network (GAN), which we think is uniquely suited for regression-based ML approaches for the immune system, to discover associations between the epitope and the variable chain features. This approach requires a large data stream of antigen and cognate antibody sequences, which until recently was difficult to obtain. A recently described single B-cell receptor (BCR) specific tagging method coupled with single cell deep sequencing (“linking B cell receptor to antigen specificity through sequencing” or LIBRA-seq) can rapidly isolate and sequence the BCR variable chain coding regions that can bind with high selectivity to antigenic epitopes. Towards the specific project goals, in Task 1, LIBRA-seq will be used to rapidly identify and generate candidate immunoglobulin coding sequences in response to specific linear and nonlinear epitopes (against controls), chosen through computational/molecular modeling and prioritized with SARS-CoV-2 Spike protein epitopes (but not restricted to these), injected into a mouse model, to generate large training sets; in Task 2, these training sets, along with other data sets already available in public databases, will generate a series of structural features (described above), which will be used to train the GAN; in Task 3, the predicted epitope-antibody interactions will be validated by direct experiments with synthetic antibody and phage-display systems. Thus, the proposed strategy combines foundational principles in evolutionary biology, genomics, structural chemistry, and computer science to the solution of a general biological engineering problem. Results from this project are expected to lay the foundations for a rigorously tested and fully automated machine-learning system that could rapidly generate synthetic antibody candidates from the structure of a novel virus protein, which can enhance the rapid response ability against a future pandemic. The ability to develop targeted antibody therapy against non-infectious or chronic diseases, and on the production of antibody-based industrial enzymes, will also be dramatically enhanced if this project were to be successful. The team: The team-leads of this multi-institutional research project comprise a computer scientist, a protein crystallographer, an immunologist, and a molecular biologist.

**Public Health Relevance Statement**

The readiness for a future viral pandemic will critically depend on being able to accurately predict, entirely using computers, neutralizing antibody structures directly from the virus’s protein structure. Vaccines and therapeutics can then be manufactured and tested with unprecedented rapidity. We propose to make this goal a reality by engaging the advantages of single-cell genomic deep sequencing and the latest advances in deep machine learning.

**NIH spending category**

Bioengineering; Biotechnology; Coronaviruses; Coronaviruses Therapeutics and Interventions; Data Science; Emerging Infectious Diseases; Genetics; Human Genome; Immunization; Infectious Diseases; Machine Learning and Artificial Intelligence

**Related publications:**

`o` Sereshki S, Lonardi S. Predicting Differentially Methylated Cytosines in TET and DNMT3 Knockout Mutants via a Large Language Model. Preprint. bioRxiv. 2024;2024.05.02.592257. Published 2024 Sep 4. doi:10.1101/2024.05.02.592257
`o` Singh P, Lonardi S, Liang Q, et al. Babesia duncani multi-omics identifies virulence factors and drug targets. Nat Microbiol. 2023;8(5):845-859. doi:10.1038/s41564-023-01360-8
`o` Ray A. Machine learning in postgenomic biology and personalized medicine. Wiley Interdiscip Rev Data Min Knowl Discov. 2022;12(2):e1451. doi:10.1002/widm.1451

